# DNA supercoiling-mediated collective behavior of co-transcribing RNA polymerases

**DOI:** 10.1101/2021.03.04.433986

**Authors:** Shubham Tripathi, Sumitabha Brahmachari, José N. Onuchic, Herbert Levine

## Abstract

Multiple RNA polymerases (RNAPs) transcribing a gene have been known to exhibit collective group behavior, causing the transcription elongation rate to increase with the rate of transcription initiation. Such behavior has long been believed to be driven by a physical interaction or “push” between closely spaced RNAPs. However, recent studies have posited that RNAPs separated by longer distances may cooperate via the DNA segment under transcription. Here, we present a theoretical model incorporating the mechanical coupling between RNAP translocation and the torsional response of supercoiled DNA. Using stochastic simulations, we demonstrate long-range cooperation between co-transcribing RNAPs mediated by DNA supercoiling. We find that inhibiting transcription initiation can slow down the already recruited RNAPs, in agreement with recent experimental observations, and predict that the average transcription elongation rate varies non-monotonically with the rate of transcription initiation. We further show that while RNAPs transcribing neighboring genes oriented in tandem can cooperate, those transcribing genes in divergent or convergent orientations can act antagonistically, and that such behavior holds over a large range of intergenic separations. Our model makes testable predictions, revealing how the mechanical interplay between RNAPs and the DNA they transcribe can govern a key cellular process.

## INTRODUCTION

Genomic DNA is double-stranded, the two strands wrapped helically around one another. The topology of DNA imposes a constraint on the movement of RNA polymerases (RNAPs) along the DNA during transcription, first conceptualized in the twin-domain model [1]. The model postulated that transcription would result in the overtwisting of the DNA downstream from the RNAP (positive supercoiling) and undertwisting of the DNA upstream from the RNAP (negative supercoiling). Recent experimental advances have implicated transcription-associated DNA supercoiling in transcriptional bursting [2], control of transcription elongation [3], and in the formation of chromosomal domains in bacteria [4, 5]. Simultaneously, advancements in single-molecule experiments have shed light on how molecular motors like RNAPs respond to mechanical interventions including DNA stretching and twisting [6]. Together, the experimental advances have resulted in both a need and an opportunity for the development of a theoretical framework of the transcription-supercoiling interplay [7–10] that can help with the analysis of the existing experimental data and make testable predictions to guide future study design.

Transcription in prokaryotes involves two distinct steps— transcription initiation and transcription elongation. During transcription initiation, an RNAP is recruited to the gene promoter. The two DNA strands are then separated by the RNAP to form a transcription bubble [11]. During transcription elongation, the bubble translocates along the DNA, with the RNAP tracking the DNA helical groove. Previous studies have suggested that two RNAPs co-transcribing a gene can cooperate [7, 12], causing the rate of transcription elongation to increase with the rate of transcription initiation [13]. However, it remains unclear if physical proximity between RNAPs is essential for them to cooperate. Further, whether the transcription elongation rate depends on the initiation rate across the physiological range of gene expression levels remains an unanswered question.

Here, we build upon the known physical properties of DNA as a polymer and upon other experimental observations to describe a theoretical model of the DNA supercoiling-transcription interplay. We use the model to probe the velocity profiles of individual RNAPs and to show that RNAP translocation-DNA supercoiling dynamics can lead to the emergence of a collective behavior regime wherein co-transcribing RNAPs can cooperate, resulting in an increase in the overall rate of transcription elongation. The proposed mechanism of driving collective RNAP behavior is in agreement with recent experimental findings [3]. We find that DNA supercoiling can drive coupling between RNAPs separated by long distances, including those transcribing neighboring genes, in contrast to the physical “push” between closely spaced RNAPs which has long been posited [12]. We further show that the interaction between RNAPs transcribing neighboring genes can be cooperative or antagonistic, depending on the relative orientation of the neighbors. Finally, we discuss the implications of the mechanical control of transcription elongation for the RNA production rates.

## MATERIALS AND METHODS

### A theoretical model of DNA supercoiling-transcription elongation interplay

As mentioned earlier, during transcription elongation, the transcription bubble translocates along the DNA, requiring the RNAP to track the DNA helical groove. This can be accomplished via rotation of the RNAP around the axis of the DNA double helix. However, if the rotational viscous drag on the RNAP complex, which includes the nascent RNA and the translation machinery (in prokaryotes), is too high, the DNA at the RNAP site must be rotated for the RNAP to move forward. Since the DNA in bacteria is torsionally constrained by DNA-binding proteins [14, 15], this results in supercoiling of the DNA upstream and downstream from the RNAP. The resultant DNA restoring torque must be overcome by the RNAP to continue translocating.

To model the RNAP translocation-DNA supercoiling interplay, we consider RNAP translocation over a distance *x* nm during which a total rotational angle of *ω*_0_*x* accumulates. Here, *ω*_0_ = 1.85 nm^−1^ is the natural linking number density in unstressed double-stranded DNA [16]. This accumulated angle is divided between the rotation angle of the RNAP (*θ*) and the DNA rotation at the RNAP site, or DNA twist (*ϕ*) (Fig. 1 a). We obtain the linking number constraint

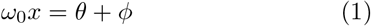

**FIG. 1.**
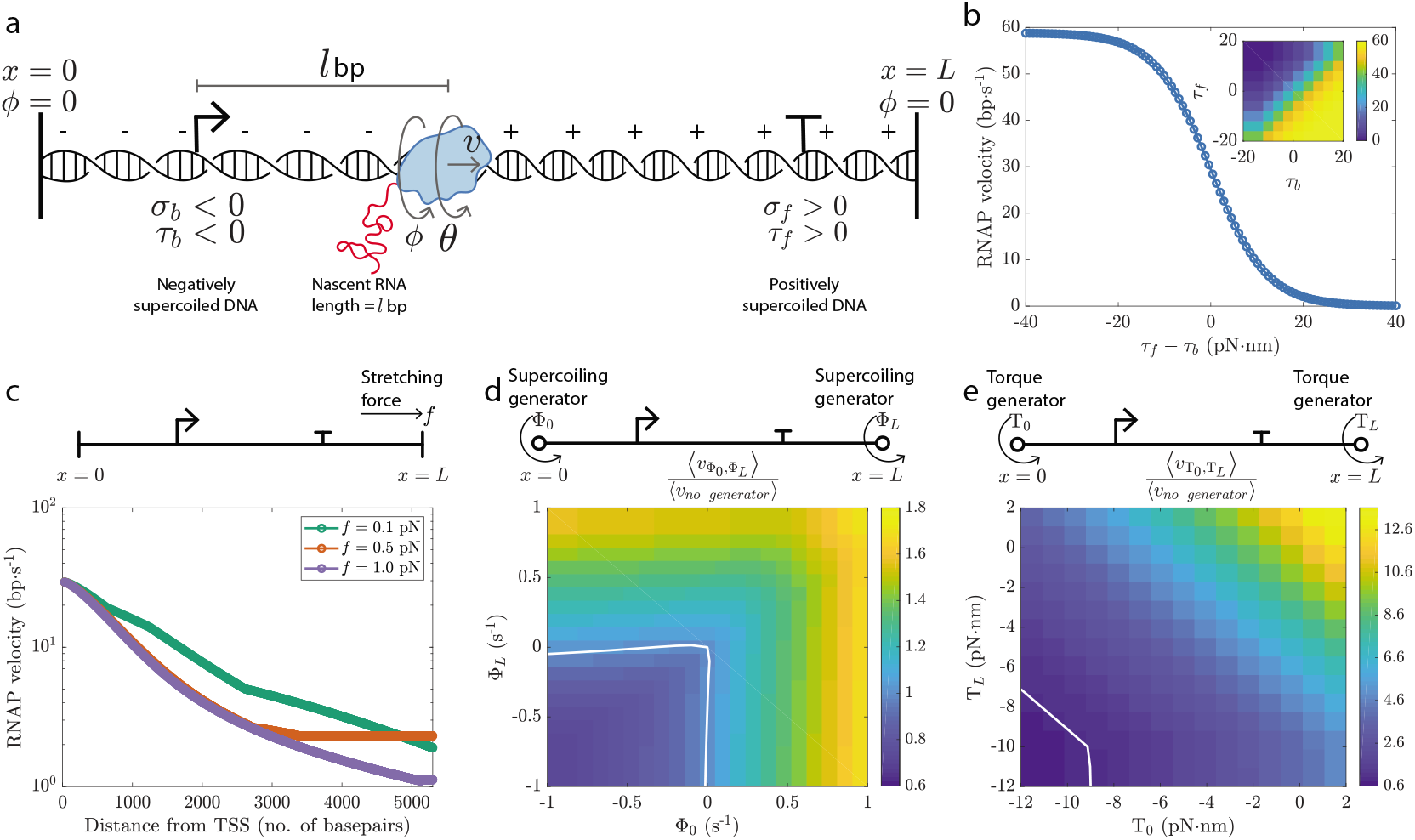
Mechanical coupling between RNAP translocation and the DNA torsional response. **a** RNAP translocation on a torsionally constrained DNA segment is accompanied by undertwisting of the DNA upstream (*σ_b_* < 0) and overtwisting of the DNA downstream (*σ_f_* > 0). The supercoiled DNA, in turn, applies a restoring torque on the RNAP (*τ_b_* and *τ_f_*). **b** In agreement with experimental data [6], we used a sigmoid curve (Eq. 3) to model the RNAP velocity-DNA restoring torque dependence. **c** RNAP velocity decreases as the RNAP moves away from the transcription start site (TSS) and is dependent on the force the DNA segment is under. **d** Ratio of the average RNAP velocity in the presence of different supercoiling generators (Eq. S4 and Eq. S5) to the average RNAP velocity in the absence of any supercoiling generators. **e** Ratio of the average RNAP velocity in the presence of different torque generators (Eq. S6 and Eq. S7) to the average RNAP velocity in the absence of any torque generators. In **d** and **e**, the white line demarcates the region where the average RNAP velocity is lower in the presence of the generators from the region where the presence of generators increases the average RNAP velocity. The average RNAP velocity is defined as the gene length divided by the total time taken by the RNAP to transcribe the gene.

The trade-off between *θ* and *ϕ* during RNAP translocation was modeled using the torque-balance equation put forth by Sevier and Levine [9]:

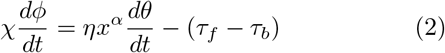

Here, *χ* is the DNA twist mobility. The first term on the right hand side of Eq. 2 describes the rotational viscous drag on the RNAP complex which grows with an increase in the nascent RNA length (equal to *x*), the growth dictated by the exponent *α*. This drag is also dependent on the coefficient of friction *η* and the rotational velocity of the RNAP complex 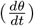. The different parameters were chosen to be within the biophysical range based on experimental data (Appendix Sec. 1).

The second term on the right hand side in Eq. 2 describes the net DNA restoring torque, equal to the difference between the restoring torque applied by the DNA segments downstream and upstream from the RNAP (*τ_f_* and *τ_b_*, respectively). The DNA restoring torque is a function of the excess linking number density, or the DNA supercoiling density *σ*, and was calculated as described previously [17, 18] (shown in Fig. S1 and Fig. S2). The restoring torque depends linearly on *σ* in regimes where supercoiling increases the DNA twist. Over the ranges of *σ* wherein increased DNA buckling or DNA melting can screen the twist, the restoring torque remains constant. These ranges of *σ* values correspond to the phase-coexistence of twisted-stretched and plectonemically buckled or melted DNA.

Experiments have shown that the RNAP velocity 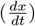 can also depend on the DNA torsional stress. Based on experimental observations [6], we used a sigmoid curve to model this dependence:

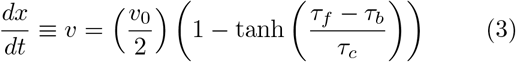

Here, *v_0_* = 20 nm·s^−1^ ≈ 60 bp·s^−1^ is the maximum RNAP velocity and RNAPs stall if *τ_f_* — *τ_b_* > *τ_c_* = 12 pN·nm. Thus, both positive torque downstream and negative torque upstream can stall an RNAP. If the net DNA restoring torque is negative (*τ_f_* < *τ_b_*), the DNA torsional response does not impede RNAP movement since in this scenario, RNAP translocation will twist the DNA to a more relaxed configuration. If the net restoring torque is positive (*τ_f_* > *τ_b_*), the DNA torsional response hinders RNAP movement since now the RNAP must further increase the DNA torsional stress in order to translocate.

We used the theoretical framework described by Eq. 1–3 to simulate the behavior of a single RNAP under different mechanical interventions and to probe how multiple RNAPs co-transcribing the same gene or neighboring genes will interact.

## RESULTS

### Translocation profile of a single RNAP

Starting with a single RNAP at the transcription start site, the dynamical system defined by Eq. 2 and Eq. 3 can be integrated numerically, under the linking number constraint in Eq. 1, to obtain the deterministic velocity profile of the RNAP. For a single RNAP transcribing a gene, the velocity decreases as the RNAP moves along the gene (Fig. 1 c). This is because while RNAP rotation (*θ*) is preferred over DNA twisting by the RNAP (*ϕ*) at the start of transcription given the low rotational drag for a short nascent RNA, the RNAP increasingly twists the DNA as transcription proceeds and the nascent RNA elongates. In response, the net restoring torque on the RNAP grows, slowing it down. This behavior is shown in detail in Fig. S4.

Note that the kinks in the velocity profile of a single RNAP seen in Fig. 1 c arise at supercoiling densities where the DNA torsional response switches from one regime to another (*e.g*., onset of phase-coexistence of stretched-twisted and plectonemically buckled DNA; Fig. S5) [17]. In experiments, the predicted kinks in the velocity profile are likely to be smoothed out due to thermal fluctuations. Since the torsional response of twisted DNA depends on the force the DNA segment is under [17, 18], the velocity profile of an RNAP is also force-dependent. At a low force (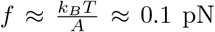, where *A* = 50 nm is the DNA persistence length and *k_B_T* = 4.1 pN·nm at room temperature), the DNA restoring torque for a given supercoiling density is weaker. Consequently, the average RNAP velocity is higher at such low forces and decreases as the force is increased (Fig. 1 c).

In Fig. 1 c, the DNA twist at the boundaries of the genomic segment is kept fixed (referred to as clamped DNA), mimicking the torsional constraint imposed by supercoiling diffusion barriers. However, in many scenarios, twist can be injected into a genomic section from the boundaries by an external agent or process. In single-molecule studies, magnetic tweezers are used to characterize the DNA’s response to twist injection. In cells, enzymes such as gyrases and topoisomerases inject negative and positive supercoiling into the DNA, respectively [19]. Both DNA transcription and replication machineries [20] can inject supercoiling into the neighboring DNA. We analyzed the RNAP velocity in the presence of supercoiling generators (Fig. 1 d), which inject supercoiling at a constant rate, and torque generators (Fig. 1 e), which inject supercoiling until the restoring torque in the DNA segment reaches a constant value. We found that generators that add negative supercoiling downstream or positive supercoiling upstream of the RNAP increase the RNAP velocity by cancelling out the RNAP-generated supercoiling. In contrast, generators that add positive supercoiling downstream or negative supercoiling upstream of the RNAP add to the RNAP-generated supercoiling, slowing the RNAP down. Together, Fig. 1 c-e show that both DNA stretching and twisting can affect RNAP translocation. The case when the DNA twisting comes from the fellow RNAPs is described next.

### Emergence of collective RNAP behavior

We incorporated the stochastic recruitment of RNAPs to the transcription start site (at a rate *k_on_*) and stochastic supercoiling relaxation events (at a rate *k_relax_*) into our model (Fig. 2 a), and explored how multiple RNAPs transcribing a gene can interact. Since a transcribing RNAP injects negative supercoiling into the upstream DNA and positive supercoiling into the downstream DNA, multiple RNAPs transcribing a gene at the same time become mechanically coupled. In particular, we found that an already transcribing RNAP speeds up if more RNAPs are subsequently recruited to the same transcription start site (Fig. 2 b). This speed-up is supercoiling-mediated, and disappears if the DNA segment is torsionally unconstrained with freely-rotating ends and no supercoiling accumulation (referred to as free DNA) or if *k_relax_* is very high (where any RNAP-generated supercoiling is quickly relaxed). Such behavior has recently been reported for *Escherichia coli* [3].

**FIG. 2.**
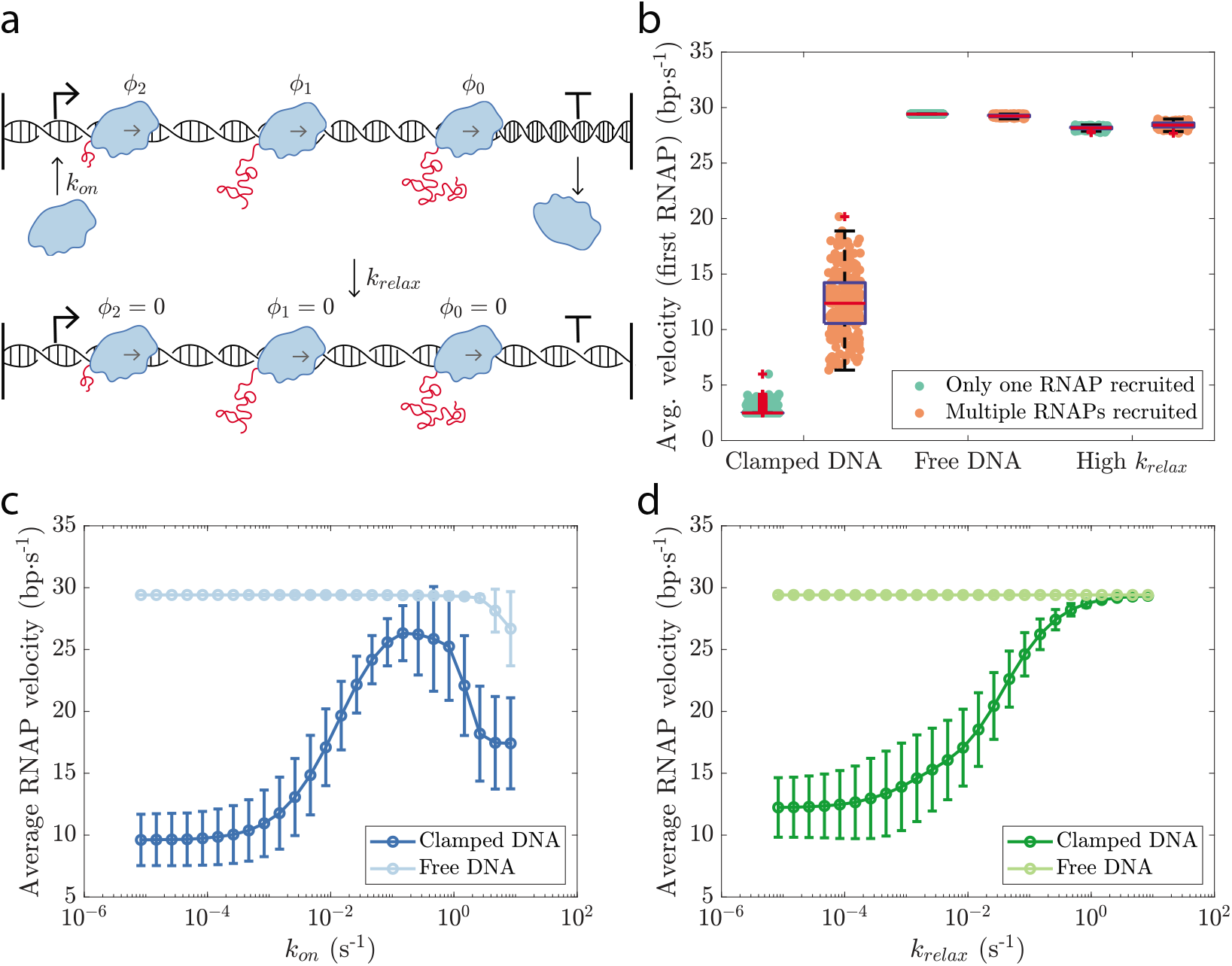
Emergence of DNA supercoiling-mediated collective behavior in co-transcribing RNAPs. **a** In our simulation setup, RNAPs are recruited to the transcription start site at a rate *k_on_* and the supercoiling throughout the genomic segment is relaxed at a rate *k_relax_*. **b** When the DNA segment is torsionally constrained (clamped DNA), the velocity of the first RNAP is higher if more RNAPs are subsequently recruited to the same gene. Fig. S6 demonstrates this behavior in greater mechanistic detail. The effect disappears if there is no supercoiling accumulation (torsionally unconstrained or free DNA) or if the RNAP-generated supercoiling is quickly relaxed (high *k_relax_*). In each case, the behavior for 256 independent runs is shown. **c** The average RNAP velocity varies non-monotonically with *k_on_* in the case of clamped DNA. Collective RNAP behavior, which emerges for *k_on_* > 10^−3^ s^−1^, increases the overall transcription elongation rate. However, for very high *k_on_*, RNAP velocity decreases due to overcrowding of the gene body (Fig. S7). **d** The average RNAP velocity increases monotonically with an increase in *k_relax_*. The error bars in **c** and **d** indicate the standard deviation. In agreement with previous studies [2, 21, 22], transcription in our setup occurs in bursts (Fig. S8). Overall, a gene under transcription injects negative supercoiling into the upstream DNA and positive supercoiling into the downstream DNA (Fig. S9).

To determine if the supercoiling-mediated interaction between co-transcribing RNAPs seen in Fig. 2 b can drive collective behavior, we varied *k_on_*, revealing three distinct regimes. At low *k_on_*, there is, on average, a single RNAP transcribing a gene at a time, and the average RNAP velocity is similar to the deterministic single RNAP case (Fig. 1 c). At higher *k_on_*, multiple RNAPs are present on the gene body at any instant, and these can cooperate. The positive supercoiling injected downstream by a trailing RNAP can at least partially be cancelled by the negative supercoiling injected upstream by the leading RNAP. The emergent collective behavior increases the average RNAP velocity by nearly 2.5-fold. At very high values of *k_on_*, a different type of collective behavior is observed— a traffic jam-like situation wherein the trailing RNAPs simply wait for the ones in the front to move, decreasing the average RNAP velocity. In the case of free DNA, the RNAP velocity remains largely unaffected by *k_on_*, confirming that the dependence of the average RNAP velocity on *k_on_* is driven by DNA supercoiling. A decline in RNAP velocity at very high values of *k_on_* is also observed in the case of free DNA, indicating that this regime is supercoiling-independent. The biological relevance of this traffic jam-like regime remains to be experimentally investigated.

Note that while DNA supercoiling by RNAPs can drive collective RNAP behavior, its overall effect remains to slow down transcription elongation— the average RNAP velocity at a fixed *k_on_* increases if the supercoiling is quickly relaxed (Fig. 2 d). Indeed, supercoiling relaxation by gyrases and topoisomerases in cells is essential for continued transcription, and inhibitors of these enzymes are potent antibacterial agents [23].

### RNAPs transcribing neighboring genes can become coupled

Until now, we have explored how the RNAPs co-transcribing the same gene can exhibit collective behavior. However, in the absence of barriers to supercoiling diffusion between genes, the RNAPs transcribing neighboring genes can also become mechanically coupled. Our simulations show that the effect of RNAPs transcribing a gene on those transcribing a neighboring gene depends on the relative orientation of the neighbors (Fig. 3). If the neighboring genes are in tandem (Fig. 3 a), the negative supercoiling injected into the intergenic region by the RNAPs transcribing the downstream gene can be cancelled by the positive supercoiling injected during the transcription of the upstream gene. Consequently, turning on the upstream gene speeds up transcription on the downstream gene, and vice versa. In contrast, when the neighboring genes are in a divergent (Fig. 3 b) or a convergent orientation (Fig. 3 c), the transcription of both neighbors injects the same type of supercoiling into the intergenic region (negative supercoiling in the case of divergent genes and positive supercoiling in the case of convergent genes). Therefore, in both cases, transcription on the downstream gene can be slowed by turning on the upstream gene, and vice versa.

**FIG. 3.**
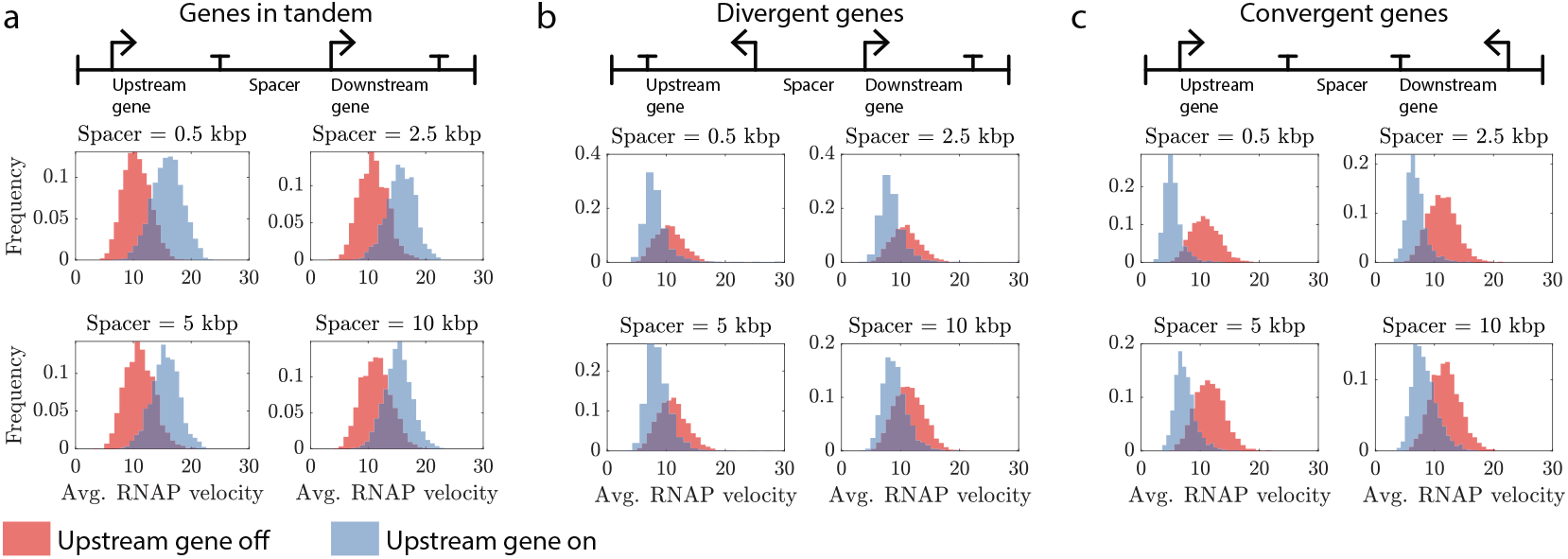
Mechanical coupling between the RNAPs transcribing neighboring genes is dependent on the relative orientation of the neighbors. Each panel shows the distribution of the average velocity of the RNAPs transcribing the downstream gene when the upstream gene is “off” (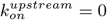) and when the upstream gene is “on” (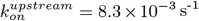). The average RNAP velocities are in units of bp·s^−1^. The histograms in each panel exhibit similar trends despite a 20-fold change in the intergenic distance. Similar behavior is observed for the reverse setup shown in Fig. S10. kbp: kilo base pairs

In agreement with our model, Kim *et al*. [3] have experimentally demonstrated, for the case of divergent neighbors, the slow down of transcription on the downstream gene if the upstream gene is highly expressed. While other studies have shown that the expression level of a gene can depend on the expression levels and the relative orientation of neighboring genes [24, 25], our model predictions concerning in tandem and convergent gene neighbors remain to be directly probed via experiments.

Note that in Fig. 3, the effect of transcription of a neighboring gene on the RNAP velocity is largely independent of the intergenic separation. Indeed, Kim *et al*. have experimentally shown that the antagonistic interaction between the RNAPs transcribing divergent genes is unaltered over a 10-fold variation in the intergenic distance [3]. In our model, this behavior emerges from the *σ*-independence of the DNA restoring torque in regimes wherein stretched-twisted DNA can coexist either with plectonemically buckled DNA or with melted DNA. We predict that in the experiments by Kim *et al*. probing divergent genes [3], stretched-twisted DNA and melted DNA co-exist in the negatively supercoiled intergenic region for the different intergenic separations.

### Effect of supercoiling-transcription interplay on RNA production

Finally, when *k_on_* is high and transcription elongation rather than transcription initiation is the rate limiting step in RNA production, DNA supercoiling-mediated processes can also alter the mean RNA production rate. Consistent with the overall repressive effect of DNA supercoiling on transcription elongation, the mean RNA production rate is higher if the RNAP-generated supercoiling is quickly relaxed (Fig. 4 a). Similarly, the mean RNA production rate can also be increased by relieving the antagonistic supercoiling being generated from the transcription of a neighboring gene as shown for the case of a convergent gene pair in Fig. 4 b. These predictions concerning the supercoiling-mediated control of RNA production rates can inform the design of synthetic circuits that exhibit predictable behaviors [24, 26].

**FIG. 4.**
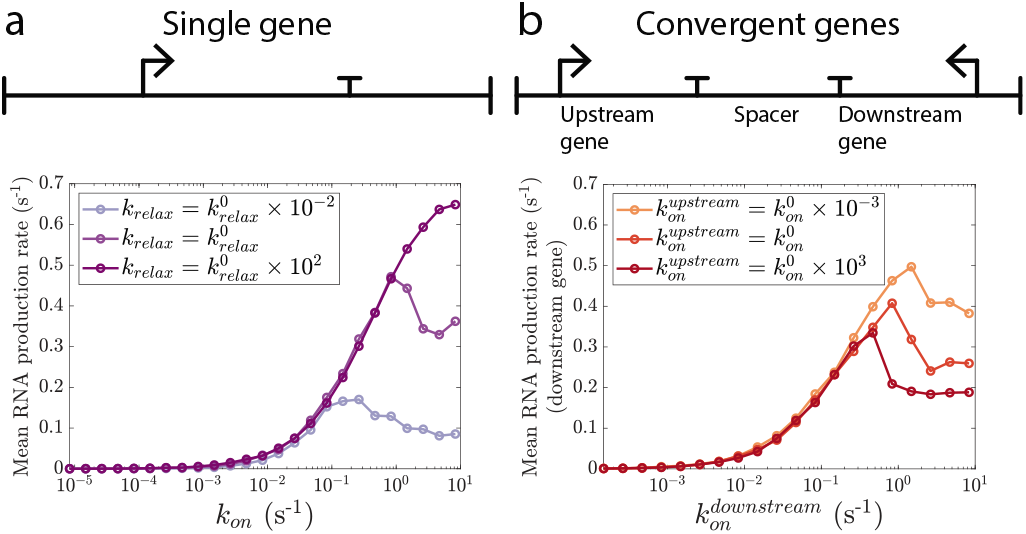
Under high transcription initiation rates, DNA supercoiling-mediated processes can alter the RNA production rates. Here, the mean RNA production rate is the number of RNAPs that finish transcribing per second, on average. **a** Rate of RNA production can be increased by quickly relaxing the RNAP-generated supercoiling. Preliminary analysis suggests that the supercoiling relaxation rate can also modulate the shape of the response to a gene inducer (Fig. S12). **b** In a setup with convergent genes, the positive supercoiling injected by the upstream gene slows down the RNA production from the downstream gene provided transcription initiation at the downstream gene is not limiting (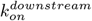 is high). The behavior in the case of in tandem and divergent neighbors is shown in Fig. S11.

## DISCUSSION

The DNA supercoiling-transcription interplay has been of interest for several decades [27], with recent experimental advances revealing the microscopic details [2–4] and a number of theoretical models posited to explain the observed behavior [7–10]. Here, we have described a framework which relies only on the mechanical behavior of DNA as a twistable, semi-flexible polymer to show that collective group behavior can emerge at the molecular level from purely mechanical processes. The model captures how transcription-mediated supercoiling can lead to coupling between the RNAPs transcribing neighboring genes, with crucial implications for the transcriptional response of genes. The framework presented here relies only upon the physical properties of DNA to make multiple testable predictions regarding RNAP dynamics, and can easily be generalized to case of other DNA-twisting motors such as DNA polymerases [20].

In agreement with experimental findings [3], simulations of our theoretical model show that cooperation between co-transcribing RNAPs can be driven by the mechanical response of the DNA segment under transcription, and does not require physical proximity between the RNAPs. Importantly, our model predicts that the cooperation between RNAPs contributes to scaling of the transcription elongation rate with the transcription initiation rate only if the transcription initiation rate is within a range— the transcription elongation rate is independent of the transcription initiation rate at low initiation rates while the elongation rate begins to decline at very high initiation rates. Thus, the initiation rate-elongation rate scaling can vary from one promoter to another depending on the promoter strength.

Understanding how DNA supercoiling affects transcription could be key to progress in synthetic biology when it comes to designing gene constructs that exhibit predictable gene expression patterns [26]. Studies have begun incorporating DNA supercoiling considerations into the design of synthetic circuits [24]. Our model shows that the rate of RNA production can depend on topoisomerase activity— variation in topoisomerase activity from cell to cell can be a driver of heterogeneity in gene expression. Preliminary analysis (shown in Fig. S12) further suggests that the topoisomerase activity can also modulate the shape of the response to a gene inducer. Our model also predicts that for genes in convergent orientation, rate of RNA production from a gene can depend on the activation level of the neighboring gene. For gene pairs in tandem or divergent orientation, in contrast, the mean RNA production rate from a gene is largely unaffected by expression from a neighboring gene (shown in Fig. S11). With these predictions, our model can be helpful in guiding the design of synthetic circuits that eliminate the undesired effects of DNA supercoiling on circuit response, and can harness supercoiling-related processes to one’s advantage.

Note that the present model assumes that the transcription initiation rate is independent of the supercoiling density at the promoter site. Others have proposed that the rate of transcription initiation rate can vary with the supercoiling density in the promoter region— a sigmoidal dependence in [28], a linear dependence in [8], and a more complex dependence based on the free energy of transcription bubble-formation in [10]. However, experimental data on this front remains scant. Extending the present model to include the supercoiling-dependence of transcription initiation on the basis of new experimental observations would make the model more useful for predicting the effect of supercoiling dynamics on the response of gene circuits to varying inducer concentrations. Such an extension of the model would also be helpful in understanding the role of DNA supercoiling in mediating the bacterial response to stress or nutrient deprivation [29, 30].

The present model is limited to the DNA supercoiling-transcription interplay in prokaryotes. In eukaryotes, the genomic DNA is wrapped around histones which change the linking number of DNA by introducing writhe [31]. Transcription elongation in eukaryotes proceeds with the expulsion of histones which may be facilitated by the torsional stress introduced by an RNAP [32]. Moreover, histones can serve as a buffer for the positive twist injected into the upstream DNA [33]. Incorporating these effects will be key to understanding the supercoiling-transcription interplay in eukaryotes as well as the role of supercoiling in chromatin organization [34].

## Appendix Methods

### 1. RNAP dynamics

From Eq. 1 and Eq. 2, we have

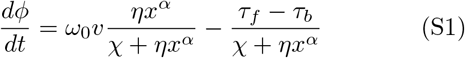

Eq. S1 and Eq. 3 were integrated numerically to simulate the dynamics of an RNAP. The different model parameters were estimated from experimental observations as described in the subsequent paragraphs. Note that the upstream and downstream DNA restoring torques (*τ_b_* and *τ_f_* in Eq. 2, Eq. 3, and Eq. S1, respectively) are defined with respect to the direction of RNAP movement—the downstream DNA segment is the DNA segment towards which the RNAP is moving (see Fig. 1 a).

In Eq. 2 and Eq. S1, *χ* is the DNA twist mobility which captures the viscous drag that will act on a DNA segment being twisted at a given rate. Approximating a DNA segment with a cylinder of radius of cross-section 1 nm that is being rotated around its long axis, we have *χ* = 8*πμ*(1 nm)^2^*C* ≈ 0.05 pN·nm·s. Here, *μ* = 17.5 Pa·s is the viscosity of the bacterial cytoplasm (BNID 108527 in [35]) and *C* = 100 nm is the experimentally reported DNA twist persistence length [17].

The model parameters *η* and *α* characterize the rotational viscous drag acting on the RNAP-nascent RNA complex. We consider three possible scenarios (illustrated in Fig. S13 a):

1. If the nascent RNA is aligned with the axis of RNAP rotation, we can approximate the rotating RNAP-nascent RNA complex with a cylinder of length equal to the nascent RNA length (*x*) and radius of cross-section 1 nm which is rotating around its long axis. In this scenario, the rotational viscous drag will equal 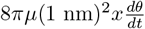. Thus, *η* ≈ 5 × 10^−4^ pN·s and *α* = 1. This scenario is more likely when the number of RNAPs simultaneously transcribing a gene is small or the nascent RNA length is small, allowing the nascent RNAs to stretch along the DNA double helix.
2. We may assume that the nascent RNA behaves like a random polymer with a persistence length *A* = 1 nm. In this scenario, the RNAP-nascent RNA complex can be treated as a sphere of radius 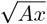 where x is the length of the nascent RNA. The rotational viscous drag on the RNAP-nascent RNA complex will then equal 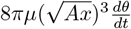, giving *η* ≈ 5 × 10^−4^ 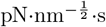 and *α* = 3/2.
3. Finally, one may approximate the RNAP-nascent RNA complex with a cylinder of length equal to the nascent RNA length (*x*) and radius of cross-section 1 nm which is rotating around an axis perpendicular to its long axis. The rotational viscous drag on the RNAP-nascent RNA complex will then equal 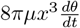, giving *η* ≈ 5 × 10^−4^ pN·nm^−2^·s and α = 3. This scenario is more likely when the density of RNAPs on the gene body is very high, causing the nascent RNAs to stretch out perpendicular to the DNA segment due to the steric hindrance from the neighboring nascent RNAs.

Note that *α* = 1 and *α* = 3 are boundary cases and that the value of *α* in experimental setups and biological systems will be somewhere in between. Further, the value of *α* will likely vary with the length of the nascent RNA, the density of RNAPs on the gene body, and the rate of recruitment of ribosomes to the nascent RNA during the co-transcriptional translation in bacteria. In the present study, we have used *α* = .3/2 since the approximation of the RNAP-nascent RNA complex as a sphere is likely to be a good assumption over a broad range of model conditions (as compared to *α* = 1 or *α* = 3 which, as described above, are boundary cases). Fig. S13 b, c show that the key model behavior is unaffected by the choice of *α*. The detailed rotational dynamics of an RNAP, however, remain uncharacterized. The rotational drag on the RNAP-nascent RNA complex is likely to include a nascent RNA length-independent contribution and a dependence on the recruitment of ribosomes to the nascent RNA during the co-transcriptional translation in prokaryotes. Both of these contributions are not included in the present study.

Finally, the results presented in this manuscript are not dependent on the form of the sigmoid function used to describe the dependence of RNAP velocity on the net DNA restoring torque. Replacing the tanh function in Eq. 3 with the logistic function, for example, will not change the model predictions.

### 2. DNA torsional response

We consider a section of the prokaryotic genome of length *L* with one or more genes. The transcription start sites are located at 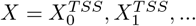, and we assume that no two gene bodies overlap. The torsional response of a DNA segment, or the restoring torque, is a function of the supercoiling density in the segment [17, 18]. Here, a DNA segment refers to any length of DNA between two barriers to supercoiling diffusion. Such barriers include the RNAPs transcribing a gene. Thus, at any instant, the genomic section is divided into segments by the *N* RNAPs present at positions {*X*_1_,*X*_2_, …,*X_N_*}. We assume that there are fixed barriers to supercoiling diffusion at both ends of the genomic section, *i.e*., at *X* = 0 and *X* = L. In single-molecule studies, these barriers could correspond to a fixed surface onto which the DNA has been immobilized or to a bead attached to the DNA end. In bacteria, the barriers would correspond to nucleoid-associated proteins which can constrain DNA supercoiling [14].

Let *ϕ_i_* and *ϕ*_*i*+1_ be the DNA rotation angles at the sites of the *i^th^* and (*i* + 1)^th^ RNAPs, respectively. Then, the supercoiling density *σ* in the DNA segment between the two RNAPs is

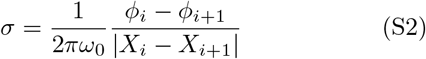

Here, we assume that the twist generated at the two ends of the DNA segment diffuses throughout the length of the segment at time scales much faster than that of RNAP dynamics. Thus, *σ* depends only on the DNA twist at the two ends of the DNA segment and is constant throughout the segment. Alternately, one can solve a transport equation for the DNA twist to probe the supercoiling dynamics at shorter time scales. Finally, we fix *ϕ*(*X* = 0) = *ϕ*(*X* = *L*) = 0 in the case of a genomic section with clamped ends. In the case of a genomic section with free ends, *ϕ*(*X* = 0) = *ϕ_N_* and *ϕ*(*X* = L) = *ϕ*_0_ (shown in Fig. S3 b).

The DNA restoring torque as a function of the supercoiling density has been described previously [17, 18] and is shown in Fig. S1 for different values of the DNA stretching force. Note that Fig. S1 shows the behavior for long DNA segments wherein plectonemes can form for *σ* > *σ_s_*. However, due to DNA’s bending stiffness [16], a short DNA segment is likely to form plectonemes only at very high supercoiling densities. To incorporate this behavior, we include a length dependence of *σ_s_* by introducing a length scale *l*_0_ below which DNA segments can accommodate higher supercoiling density before plectoneme formation can commence:

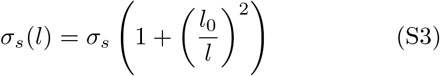

Here, *l* is the length of the DNA segment, *σ_s_* is the supercoiling density above which long DNA segments can form plectonemes (calculated in [17]) and *l*_0_ = 340 nm (or 1000 bp). The effect of the dependence of *σ_s_* on *l* on the DNA restoring torque is shown in Fig. S2.

The above description suggests that under some circumstances, plectonemes can form in the gene body. While such plectonemes may interfere with RNAP translocation due to steric hindrance, the relevance of this effect to transcription *in vivo* remains unclear. The present model does not incorporate any steric effect that plectoneme formation in the gene body may have on the translocating RNAPs.

Unless specified otherwise (see Fig. 1 c and Fig. S5, for example), the DNA restoring torque during all simulation runs was calculated under the assumption that the genomic segment is under a stretching force *f* = 1.0 pN.

### 3. Supercoiling and torque generators

The behavior of supercoiling generators was modeled using the equations

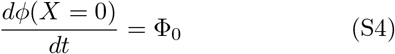

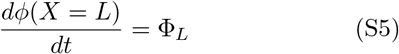

where Φ_0_ and Φ_L_ are constants. Thus, a supercoiling generator will twist the DNA at a constant rate, irrespective of the restoring torque applied by the DNA.

The behavior of torque generators was modeled using the equations

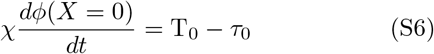

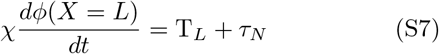

where T_0_, T_*N*_ are constants, and *τ*_0_, *τ_N_* are the restoring torques in the DNA segments shown in Fig. S3 c. Thus, a torque generator twists the DNA till the restoring torque applied by the DNA reaches a threshold (here, T_0_ or T_*N*_).

An RNAP transcribing a gene injects positive supercoiling into the downstream DNA segment and negative supercoiling into the upstream DNA segment (see Fig. S9). As the resulting net restoring torque on the RNAP grows, the RNAP velocity decreases along with the rate of supercoiling injection by the RNAP. Finally, when the net restoring torque approaches the stall torque, the RNAP stops translocating and can no longer inject supercoiling into the DNA. Thus, an RNAP transcribing a gene acts like a torque generator of the type described above.

### 4. Stochastic simulations

The stochastic simulation setup included two types of events, recruitment of RNAPs to a transcription start site at a rate *k_on_* and relaxation of supercoiling in the entire genomic section at a rate *k_relax_* (see Fig. 2 a). In simulations involving two genes, the recruitment of RNAPs to the transcription start site of each gene was independent of RNAP recruitment at the other gene. There is no premature transcription termination in our simulation setup. Once an RNAP is recruited to the transcription start site, it can fall off the DNA only when it encounters the transcription terminator. Simulations were carried out using the Gillespie algorithm [36]. The different genomic configurations used for the simulations involving one or two genes are shown in Fig. S3.

When a new RNAP is recruited to a transcription start site, we set the RNAP rotation angle *θ* = 0. The additional DNA rotation at the site of the newly recruited RNAP (*ϕ*) was chosen such that the supercoiling densities in the DNA segments on the two sides of the newly recruited RNAP are the same and equal to the supercoiling density in the DNA segment spanning the transcription start site before the recruitment of the new RNAP. During the supercoiling relaxation event, the additional DNA rotation at the site of each RNAP was set to 0 (*ϕ*_0_ = *ϕ*_1_ = … = *ϕ_N_* = 0). We also included hard-core repulsion between RNAPs so that the RNAPs do not overlap or cross one another: the trailing RNAP cannot move forward if there is another RNAP in front of it at a distance less than 15 nm. Similarly, if there is an RNAP within a distance of 15 nm from the transcription start site, no new RNAP can be recruited.

Finally, at intervals of one second, we checked if any RNAP had gone past the transcription terminator. In such a scenario, the RNAP was removed from the simulation. The average number of such events happening per second is reported as the mean RNA production rate in Fig. 4, Fig. S11, and Fig. S12.

### 5. Simulation parameters

*Fig. 2*— Panel **b**: *k_on_* = 8.3 × 10^−3^ s^−1^, *k_relax_* = 8.3 × 10^−5^ s^−1^ for the clamped DNA and free DNA cases; *k_relax_* = 0.83 s^−1^ for the “high *k_relax_*” case. Panel **c**: *k_relax_* = 8.3 × 10^−3^ s^−1^. Panel **d**: *k_on_* = 8.3 × 10^−3^ s^−1^

*Fig. 3*— In each panel, *k_on_* = 8.3 × 10^−5^ s^−1^ for the downstream gene and *k_relax_* = 8.3 × 10 ^−3^ s^−1^. When the upstream gene is “on”, 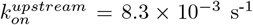, 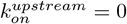 otherwise.

*Fig. 4—* Panel **a**: 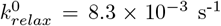. Panel **b**: 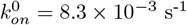.

## Supplementary Figures

**FIG. S1.**
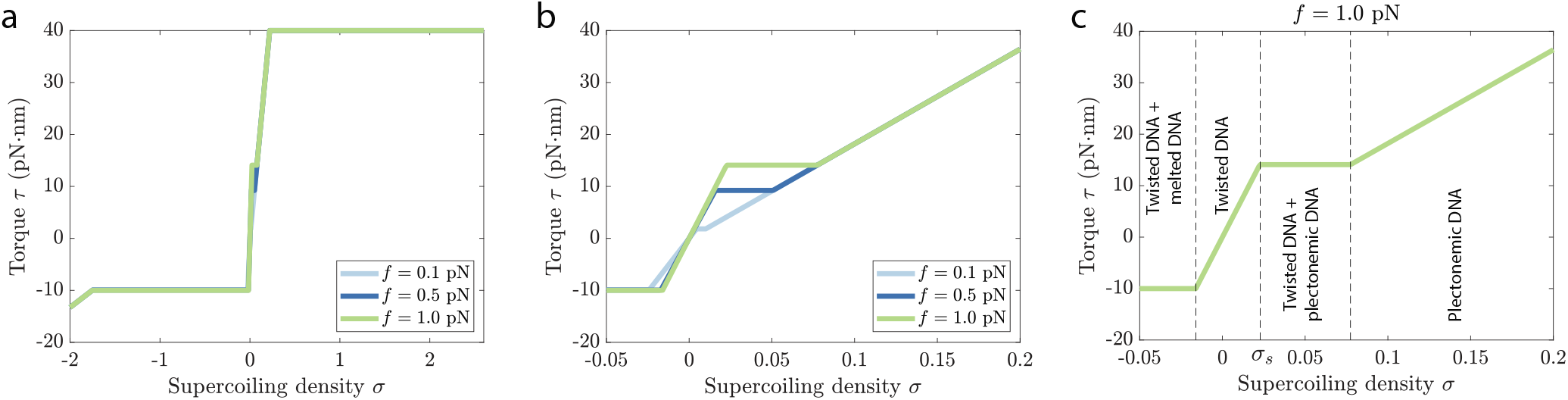
The DNA restoring torque as a function of the supercoiling density *σ*, shown for different values of the stretching force *f*. Panels **a** and **b** differ only in the range of the supercoiling densities shown on the horizontal axis. Panel **c** shows the different regimes of supercoiling densities and the DNA configuration seen in each regime. In all three panels, length of the DNA segment *l* = 10 kbp (kilo base pairs).

**FIG. S2.**
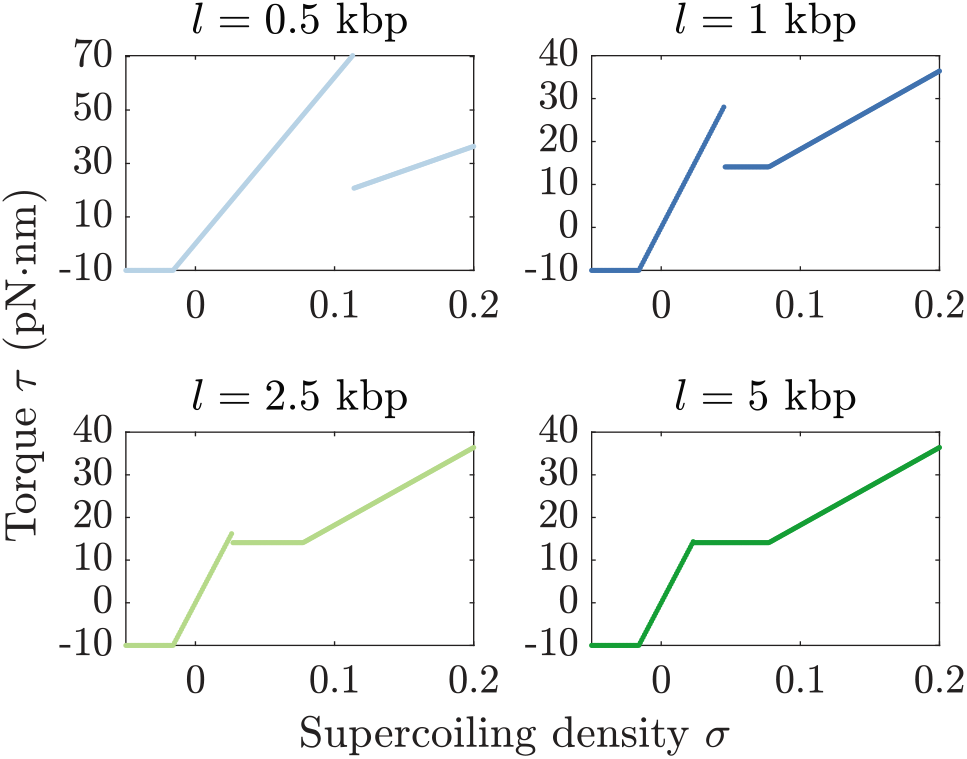
The dependence of the DNA restoring torque on the length *l* of the DNA segment for a range of supercoiling densities. Here, *f* = 1.0 pN. See Appendix Sec. 2 and Eq. S3. kbp: kilo base pairs

**FIG. S3.**
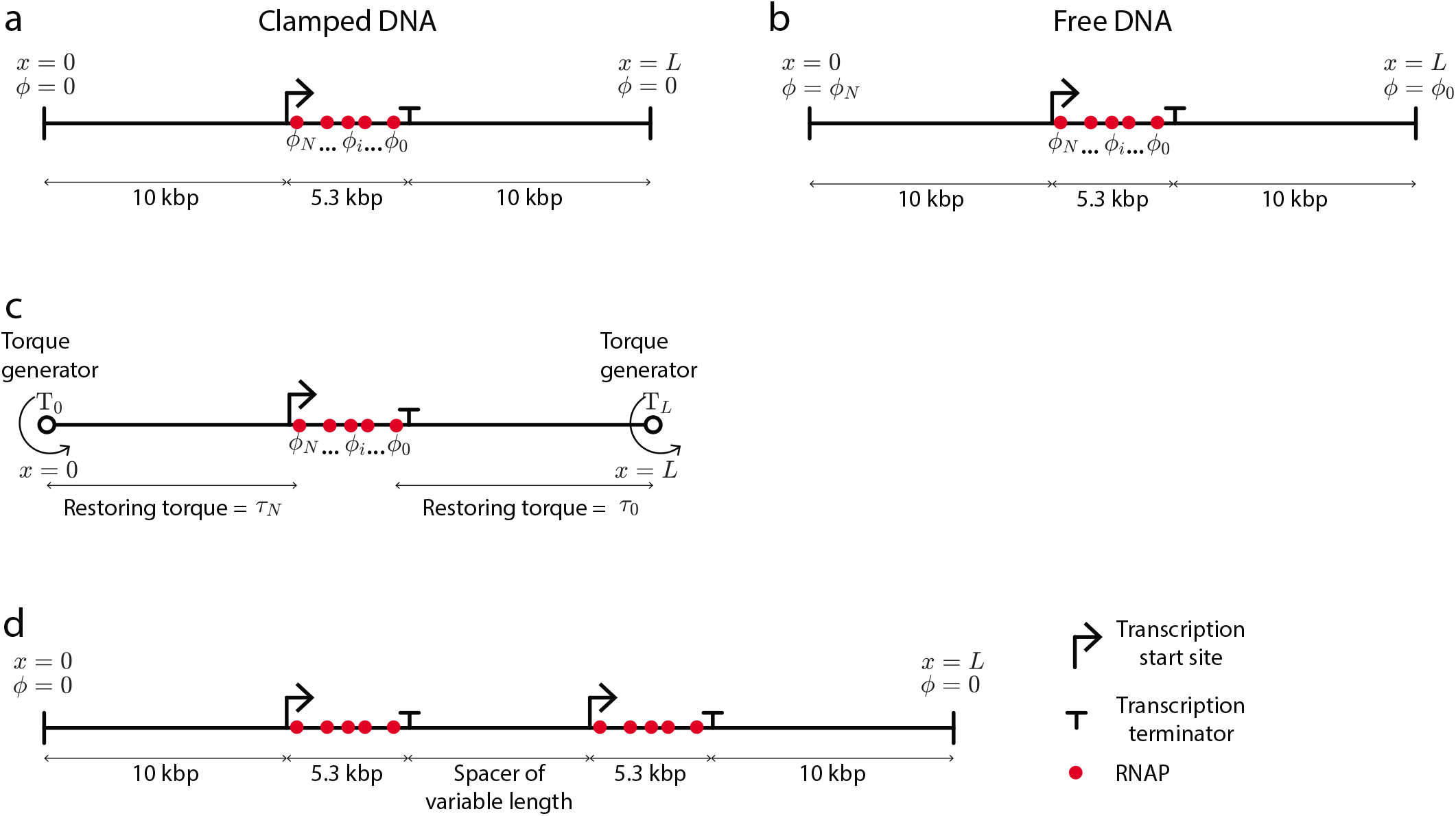
A schematic showing the different simulation setups used in the present study. **a** A single gene in a genomic segment with clamped ends. **b** A single gene in a genomic segment with free ends. **c** A single gene in a genomic segment with a torque generator at each end (see Appendix Sec. 3, and Eq. S6 and Eq. S7.). **d** Two genes in a genomic segment with clamped ends. In each panel, *ϕ* indicates the DNA rotation angle at the RNAP sites or at the ends of the genomic segment.

**FIG. S4.**
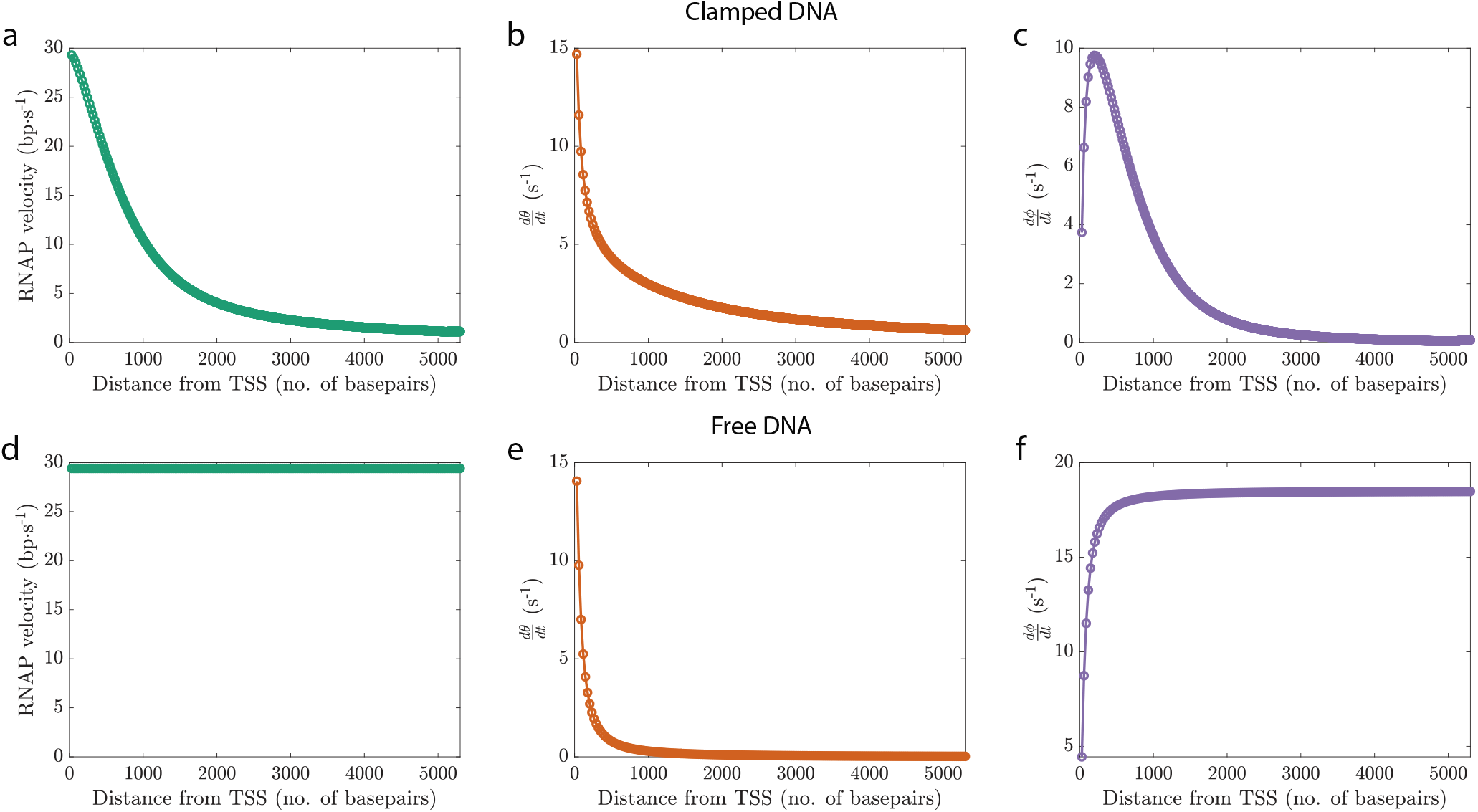
The dynamics of a single RNAP transcribing a gene as a function of the distance from the transcription start site (TSS). Since the rotational viscous drag on the RNAP-nascent RNA complex grows with an increase in the length of the nascent RNA (which equals the distance traveled by the RNAP from the TSS), the RNAP rotational velocity 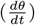 decreases with growing distance from the TSS both in the case of clamped DNA and free DNA (see panels **b** and **e**). In the case of free DNA, as the rotational drag on the RNAP-nascent RNA complex increases, the accumulating rotational angle from RNAP translocation (*ω*_0_*x*) is increasingly deposited into the DNA rotation angle at the RNAP site (*ϕ*; see panel **f**). Since the DNA has free ends, there is no DNA twist or restoring torque buildup and a constant rate of RNAP translocation is maintained (panel **d**). In the case of clamped DNA, while the rate of DNA rotation at the RNAP site 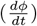 increases initially with a decline in the RNAP rotational velocity, the DNA restoring torque soon makes it impossible to maintain a DNA rotation rate high enough to keep the RNAP translocation rate constant (panel **c**). Consequently, the rate of RNAP translocation continues to decline with an increase in the distance form the TSS (panel **a**). For both clamped and free DNA, the following relation holds at each point: 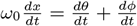.

**FIG. S5.**
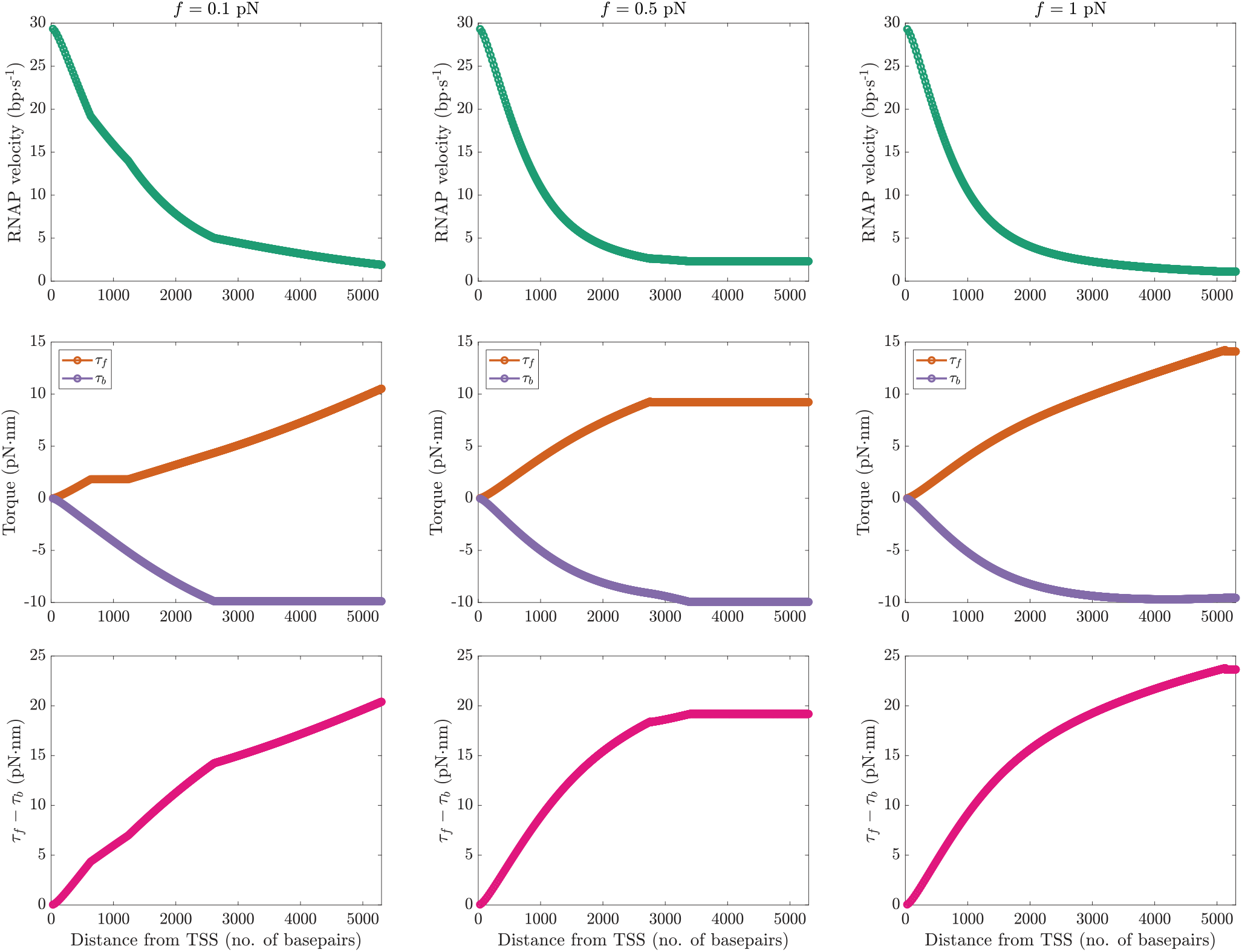
The velocity of a single RNAP and the corresponding DNA restoring torque, shown as a function of the distance from the transcription start site (TSS). Each column shows the behavior for a fixed value of the DNA stretching force (*f*). Here, *τ_b_* and *τ_f_* are the restoring torques applied by the DNA segments upstream and downstream from the translocating RNAP, respectively. Thus, the difference *τ_f_* — *τ_b_*, shown in the bottom row, is the net restoring torque on the RNAP. Note that the kinks in the RNAP velocity profiles (top row) occur at the same instants as the kinks in the torque curves (bottom two rows).

**FIG. S6.**
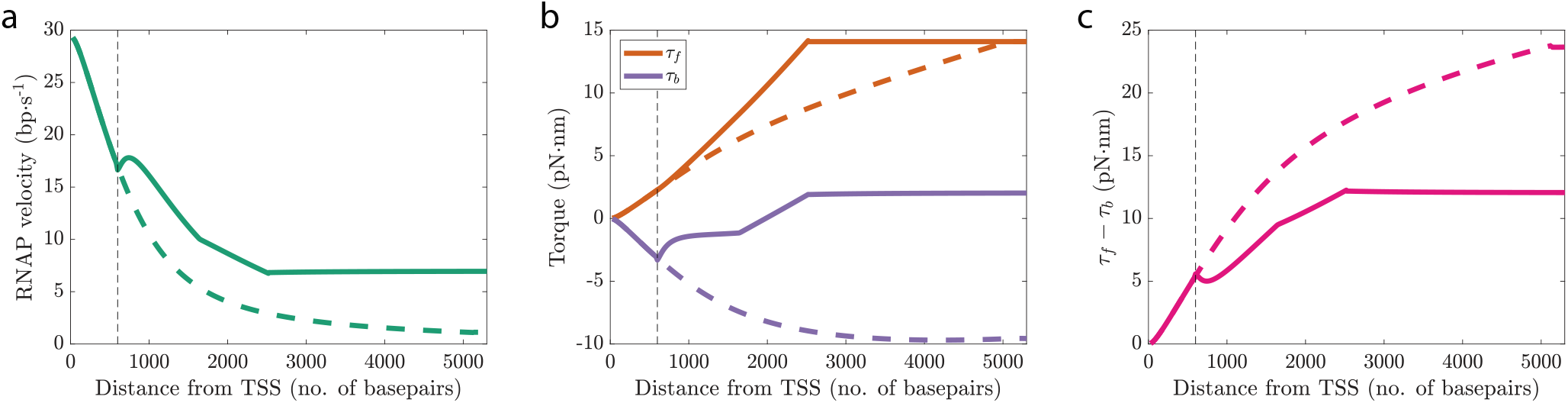
Change in the velocity profile of an RNAP when a second RNAP is recruited to the transcription start site (TSS) of the same gene. Solid curves in each panel show the behavior with respect to the first RNAP for when a second RNAP is recruited before the first RNAP has finished transcribing. The dashed curves indicate the behavior when no second RNAP is recruited. The vertical black dashed line indicates the distance of the first RNAP from the TSS when the second RNAP is recruited. When a second RNAP is recruited, the net restoring torque (*τ_f_* — *τ_b_*) acting on the first RNAP becomes lower than that in the case of a single transcribing RNAP (panel **c**). Consequently, a higher RNAP translocation rate (for the first RNAP) can be maintained (solid curve in panel **a**). Simulation parameters are the same as in Fig. 2 b.

**FIG. S7.**
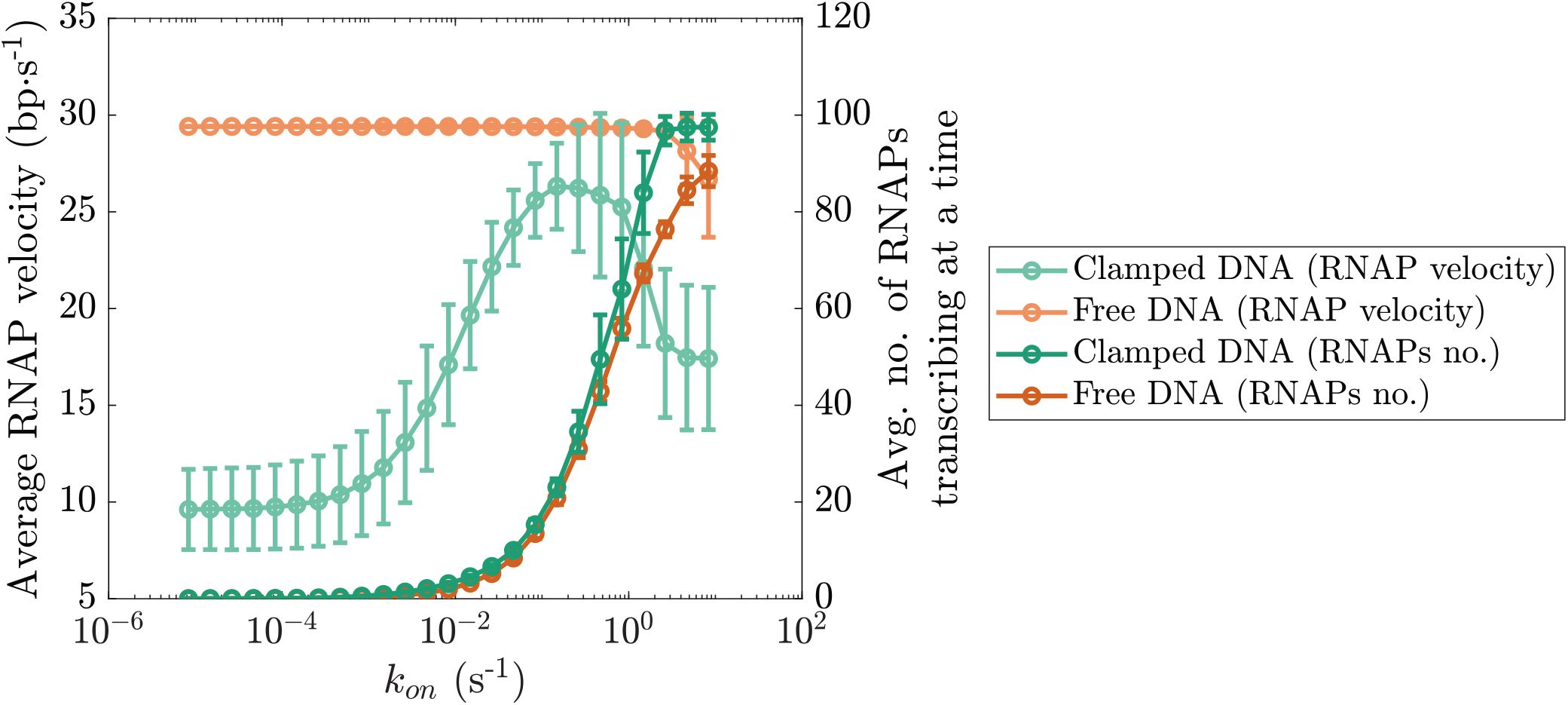
The average number of RNAPs transcribing a gene at any time as a function of *k_on_*. Simulation parameters are the same as in Fig. 2 c. The error bars indicate the standard deviation.

**FIG. S8.**
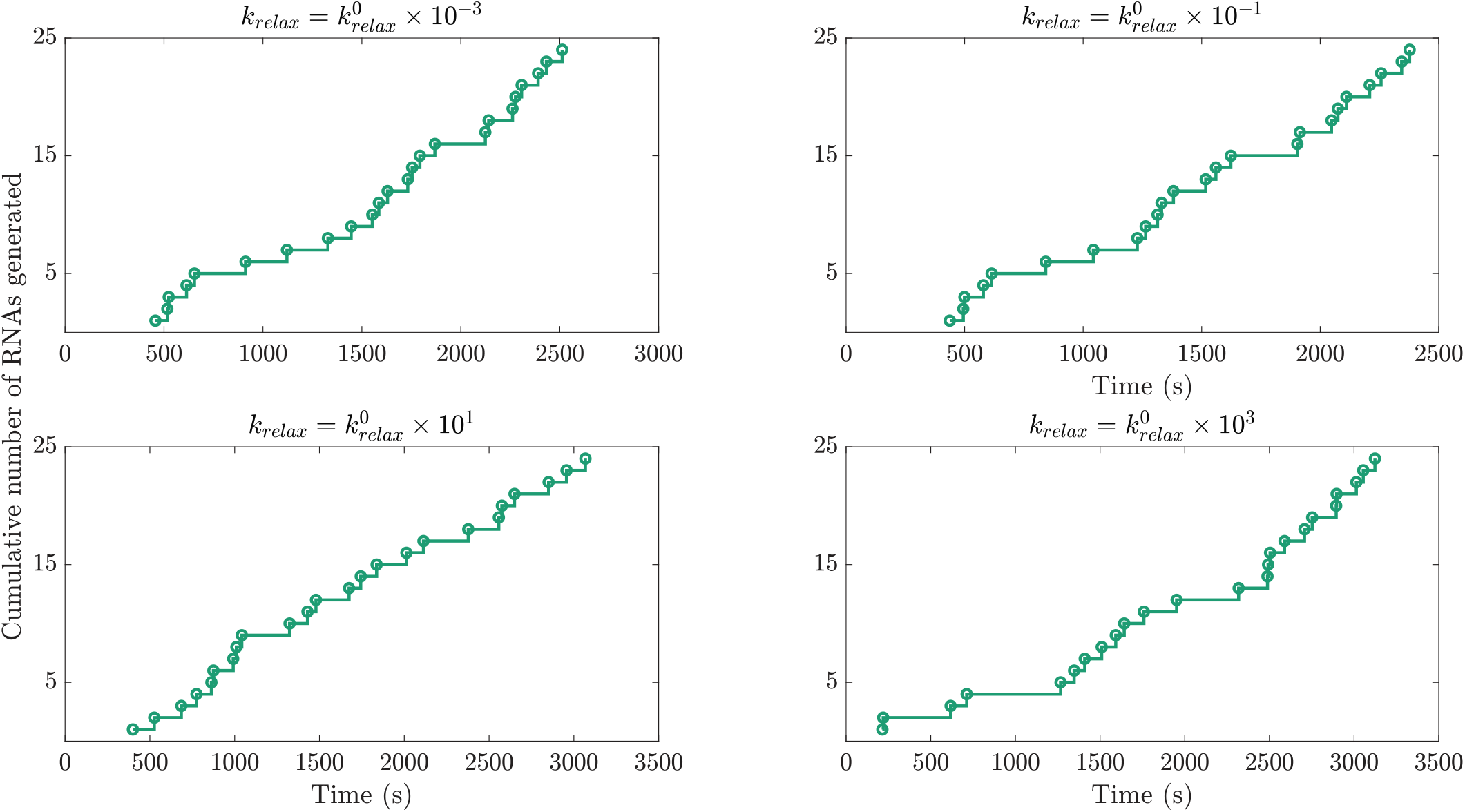
Supercoiling-mediated transcriptional bursting is observed in the case of clamped DNA. Previous studies have shown that RNAP stalling due to the DNA restoring torque buildup can result in transcriptional bursting wherein even at high induction levels, genes exhibit intervals of paused transcriptional activity [2, 21, 22]. In the case of clamped DNA, DNA twist can accumulate over time, stalling the RNAPs when the DNA restoring torque becomes too high. This results in a period of paused transcriptional activity before the DNA twist is relaxed by topoisomerase activity, jump starting RNAP translocation. In all panels, 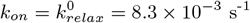.

**FIG. S9.**
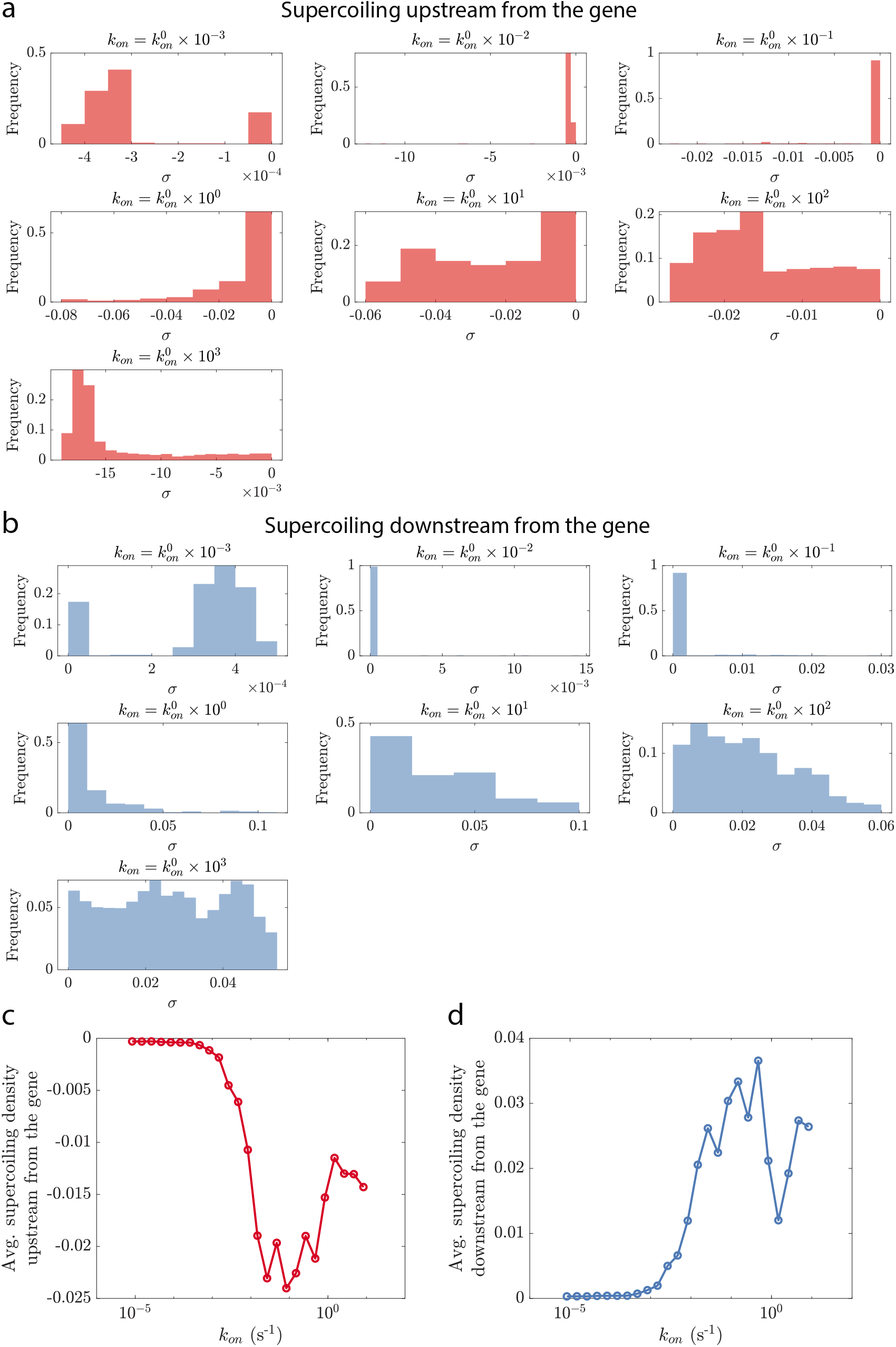
Supercoiling density in the DNA upstream from the transcription start site and downstream from the transcription terminator of a gene for different values of the RNAP recruitment rate *k_on_*. Panels **a** and **b** show the distribution of the supercoiling densities observed at different time points over a single simulation run. Panels **c** and **d** show the supercoiling density averaged over the different time points. Overall, transcription of a gene generates negative supercoiling in the upstream DNA and positive supercoiling in the downstream DNA. The magnitude of the supercoiling generated varies with *k_on_*. Here, 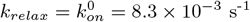.

**FIG. S10.**
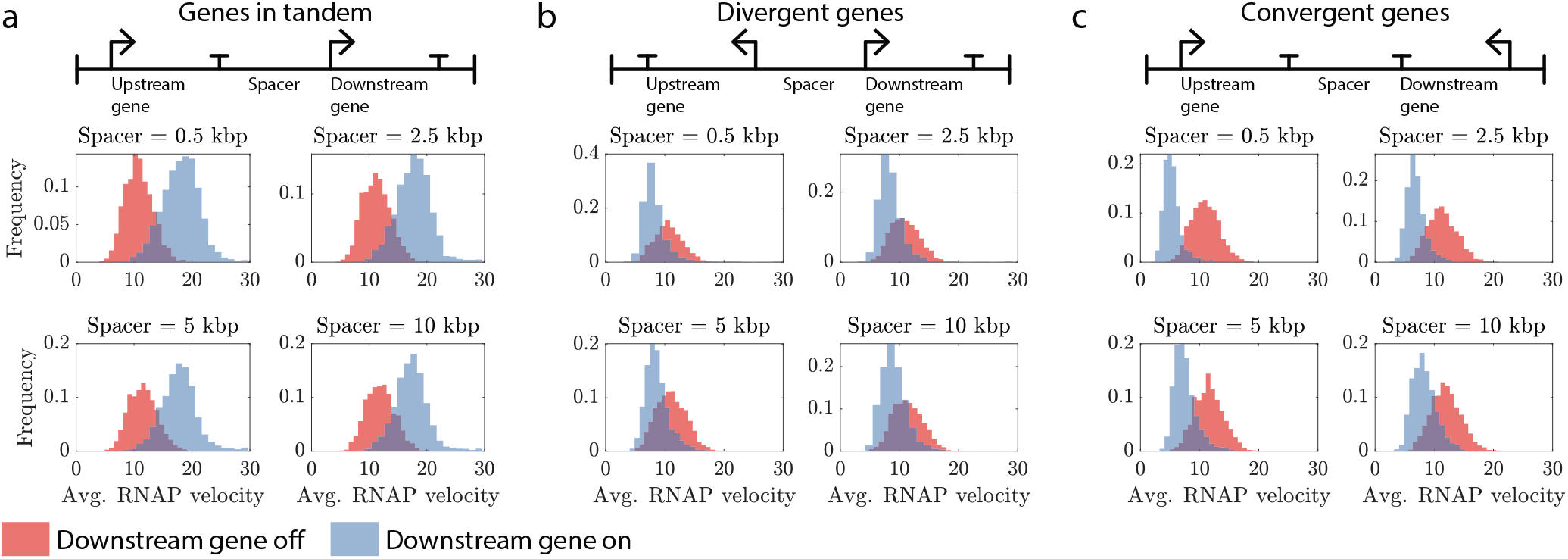
Change in the distribution of the average RNAP velocity for the upstream gene when the downstream gene is turned on. When the downstream gene is off, 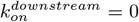. When the downstream gene is on, 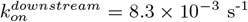. For the upstream gene, *k_on_* = 8.3 × 10^−5^ s^−1^ in each case. The behavior observed is similar to the one shown in Fig. 3. The average RNAP velocities are in units of bp·s^−1^ and *k_relax_* = 8.3 × 10^−3^ s^−1^. kbp: kilo base pairs

**FIG. S11.**
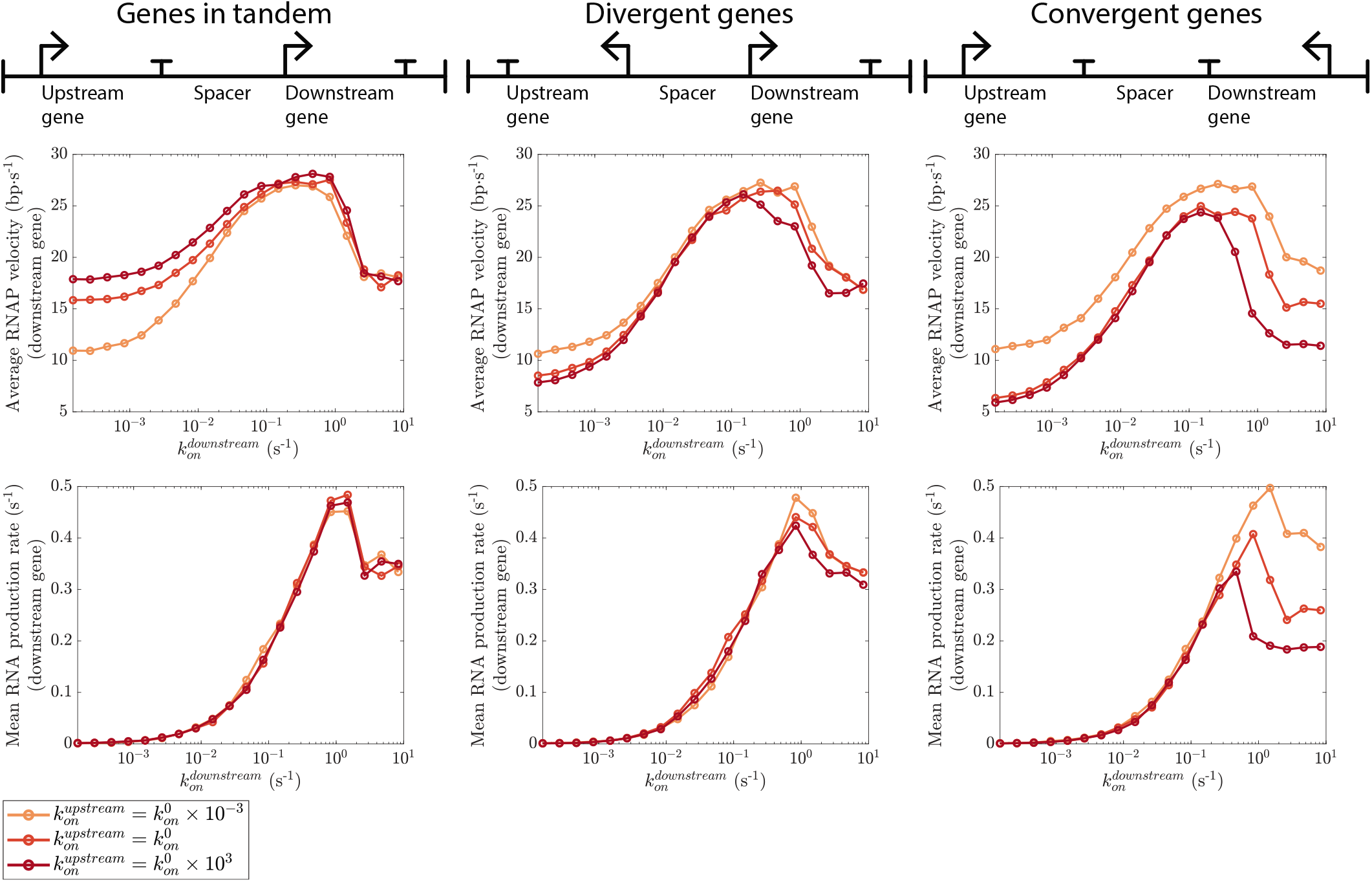
The average RNAP velocity and the mean RNA production rate for the downstream gene when the rate of RNAP recruitment to the upstream gene is varied. For in tandem and divergent gene neighbors, the average RNAP velocity for the downstream gene is affected by the RNAP recruitment rate to the upstream gene only at low values of 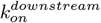. Since transcription elongation is not the rate-limiting step for RNA production at these low values of 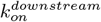, the mean RNA production rate for the downstream gene is unaffected by 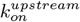. In the case of convergent genes, the average RNAP velocity for the downstream gene depends on 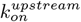 for all the values of 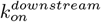 probed. Since transcription elongation is the rate-limiting step for RNA production only at high values of the RNAP recruitment rate, the mean RNA production rate for the downstream gene becomes dependent on 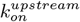 at high values of 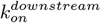. Here, 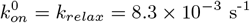.

**FIG. S12.**
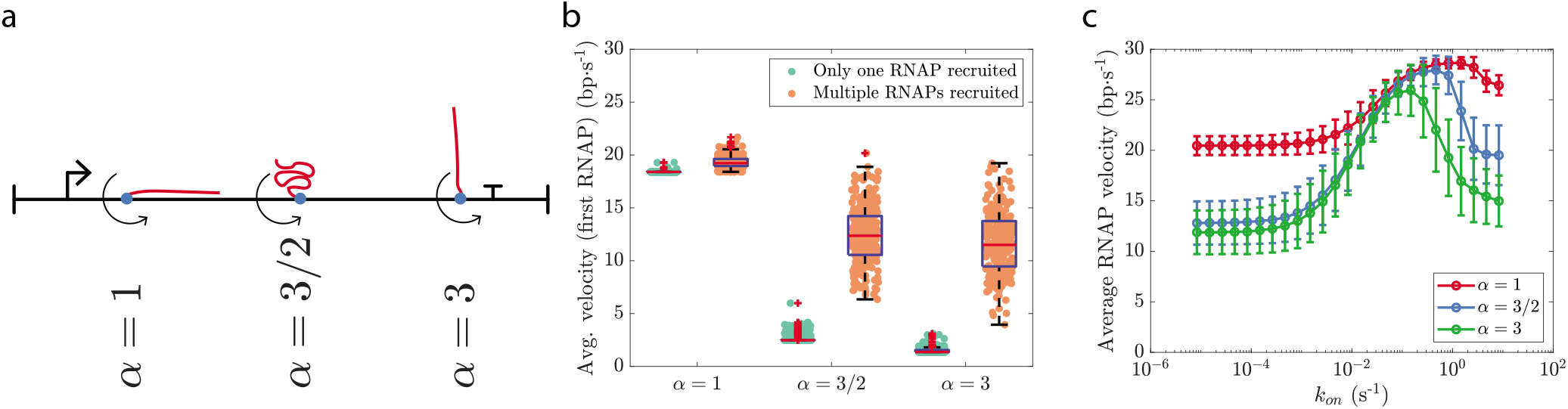
Possible effect of DNA supercoiling on the transcriptional response to an inducer. We included two additional events in our stochastic simulation setup: binding of a transcription factor to the promoter region of the gene (at a rate 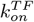) and unbinding of the transcription factor from the promoter region (at a rate 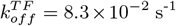). RNAPs can be recruited to the transcription start site (at a rate *k_on_* = 8.3 × 10^−2^ s^−1^) only when the transcription factor is bound to the promoter region. In the case of clamped DNA (top row), the transcription factor response curve, *i.e*., the mean RNA production rate as a function of the transcription factor recruitment rate, can depend on the rate of DNA supercoiling relaxation (*k_relax_*) (top right panel). No such dependence is observed in the case of free DNA (bottom right panel). The result suggests that the response of a cell to the induction of the activity of a transcription factor can depend on the expression levels of enzymes such as gyrases and topoisomerases. This could have important implications for the design of synthetic biology constructs with predictable response and low heterogeneity. The figure shows a preliminary result with the key assumption that the transcription factor-promoter binding rate is unaffected by the supercoiling density in the promoter region. Here, 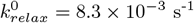.

**FIG. S13.**
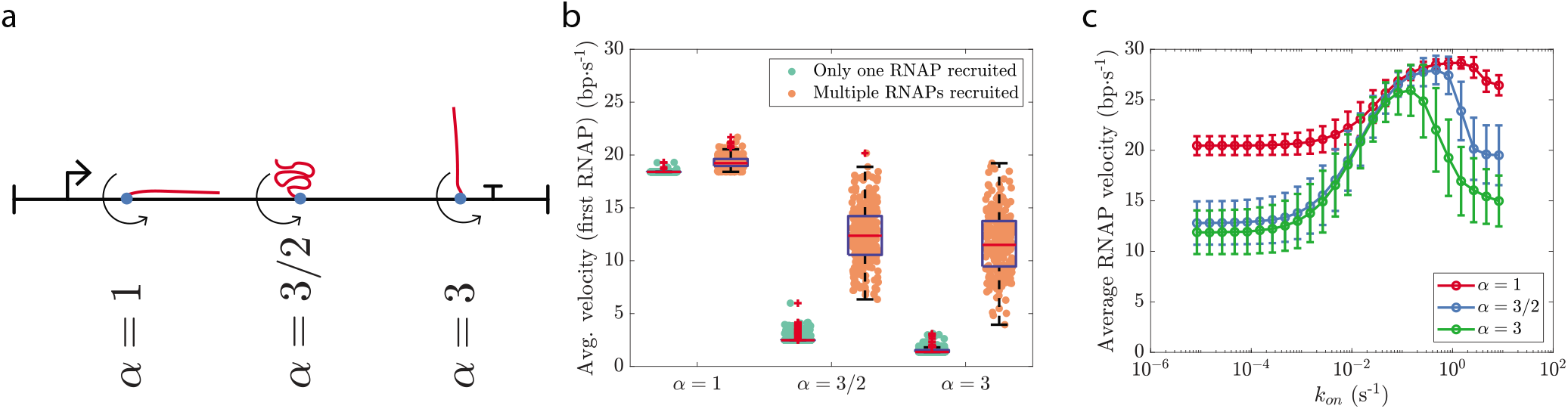
Dependence of model behavior on the parameter *α*. **a** Examples of different configurations of the nascent RNA and the corresponding values of the model parameter α. See Appendix Sec. 1 for a detailed discussion. **b** For each value of α probed, the velocity of an RNAP increases if more RNAPs are subsequently recruited to the transcription start site of the same gene. Here, *k_on_* = 8.3 × 10^−3^ s^−1^ and *k_relax_* = 8.3 × 10^−5^ s^−1^ (or 5.0 × 10^−3^ min^−1^). **c** The average RNAP velocity varies non-monotonically with the rate of RNAP recruitment to the transcription start site (*k_on_*) for the different values of *α* probed. Here, *k_relax_* = 16.6 × 10^−3^ s^−1^. Panels **b** and **c** illustrate that the key model prediction— cooperation between multiple RNAPs transcribing a gene— holds over the entire range of values the model parameter *α* can take.

## References

[1] L. F. Liu and J. C. Wang, Supercoiling of the DNA template during transcription, Proc. Natl. Acad. Sci. U.S.A. 84, 7024 (1987).

[2] S. Chong, C. Chen, H. Ge, and X. S. Xie, Mechanism of transcriptional bursting in bacteria, Cell 158, 314 (2014).

[3] S. Kim, B. Beltran, I. Irnov, and C. Jacobs-Wagner, Long-distance cooperative and antagonistic RNA polymerase dynamics via DNA supercoiling, Cell 179, 106 (2019).

[4] T. B. K. Le, M. V. Imakaev, L. A. Mirny, and M. T. Laub, High-resolution mapping of the spatial organization of a bacterial chromosome, Science 342, 731 (2013).

[5] T. B. K. Le and M. T. Laub, Transcription rate and transcript length drive formation of chromosomal interaction domain boundaries, EMBO J. 35, 1582 (2016).

[6] J. Ma, L. Bai, and M. D. Wang, Transcription under torsion, Science 340, 1580 (2013).

[7] T. Heberling, L. Davis, J. Gedeon, C. Morgan, and T. Gedeon, A mechanistic model for cooperative behavior of co-transcribing RNA polymerases, PLOS Comput. Biol. 12, 1 (2016).

[8] C. A. Brackley, J. Johnson, A. Bentivoglio, S. Corless, N. Gilbert, G. Gonnella, and D. Marenduzzo, Stochastic model of supercoiling-dependent transcription, Phys. Rev. Lett. 117, 018101 (2016).

[9] S. A. Sevier and H. Levine, Properties of gene expression and chromatin structure with mechanically regulated elongation, Nucleic Acids Res. 46, 5924 (2018).

[10] S. Meyer and G. Beslon, Torsion-mediated interaction between adjacent genes, PLOS Comput. Biol. 10, e1003785 (2014).

[11] S. Borukhov and E. Nudler, RNA polymerase: the vehicle of transcription, Trends Microbiol. 16, 126 (2008).

[12] T. T. Le and M. D. Wang, Molecular highways—navigating collisions of DNA motor proteins, J. Mol. Biol. 430, 4513 (2018).

[13] V. Epshtein and E. Nudler, Cooperation between RNA polymerase molecules in transcription elongation, Science 300, 801 (2003).

[14] S. C. Dillon and C. J. Dorman, Bacterial nucleoid-associated proteins, nucleoid structure and gene expression, Nat. Rev. Microbiol. 8, 185 (2010).

[15] F. Leng, B. Chen, and D. D. Dunlap, Dividing a supercoiled DNA molecule into two independent topological domains, Proc. Natl. Acad. Sci. U.S.A. 108, 19973 (2011).

[16] J. F. Marko, Biophysics of protein–DNA interactions and chromosome organization, Physica A 418, 126 (2015).

[17] J. F. Marko, Torque and dynamics of linking number relaxation in stretched supercoiled DNA, Phys. Rev. E 76, 021926 (2007).

[18] J. F. Marko and S. Neukirch, Competition between curls and plectonemes near the buckling transition of stretched supercoiled DNA, Phys. Rev. E 85, 011908 (2012).

[19] E. L. Zechiedrich, A. B. Khodursky, S. Bachellier, R. Schneider, D. Chen, D. M. J. Lilley, and N. R. Cozzarelli, Roles of topoisomerases in maintaining steady-state DNA supercoiling in escherichia coli, J. Biol. Chem. 275, 8103 (2000).

[20] S. A. Sevier, Mechanical properties of DNA replication, Phys. Rev. Research 2, 023280 (2020).

[21] S. A. Sevier, D. A. Kessler, and H. Levine, Mechanical bounds to transcriptional noise, Proc. Natl. Acad. Sci. U. S.A. 113, 13983 (2016).

[22] A. Klindziuk, B. Meadowcroft, and A. B. Kolomeisky, A mechanochemical model of transcriptional bursting, Biophys. J. 118, 1213 (2020).

[23] A. Maxwell, DNA gyrase as a drug target, Trends Microbiol. 5, 102 (1997).

[24] E. Yeung, A. J. Dy, K. B. Martin, A. H. Ng, D. Del Vecchio, J. L. Beck, J. J. Collins, and R. M. Murray, Biophysical constraints arising from compositional context in synthetic gene networks, Cell Syst. 5, 11 (2017).

[25] S. K. Eszterhas, E. E. Bouhassira, D. I. K. Martin, and S. Fiering, Transcriptional interference by independently regulated genes occurs in any relative arrangement of the genes and is influenced by chromosomal integration position, Mol. Cell Biol. 22, 469 (2002).

[26] S. Cardinale and A. P. Arkin, Contextualizing context for synthetic biology – identifying causes of failure of synthetic biological systems, Biotechnol. J. 7, 856 (2012).

[27] C. J. Dorman, DNA supercoiling and transcription in bacteria: a two-way street, BMC Mol. and Cell Biol. 20, 26 (2019).

[28] B. El Houdaigui, R. Forquet, T. Hindré, D. Schneider, W. Nasser, S. Reverchon, and S. Meyer, Bacterial genome architecture shapes global transcriptional regulation by DNA supercoiling, Nucleic Acids Res. 47, 5648 (2019).

[29] P. Bordes, A. Conter, V. Morales, J. Bouvier, A. Kolb, and C. Gutierrez, DNA supercoiling contributes to disconnect *σ^S^* accumulation from *σ^S^*-dependent transcription in Escherichia coli, Mol. Microbiol. 48, 561 (2003).

[30] N. Blot, R. Mavathur, M. Geertz, A. Travers, and G. Muskhelishvili, Homeostatic regulation of supercoiling sensitivity coordinates transcription of the bacterial genome, EMBO Rep. 7, 710 (2006).

[31] F. Kouzine, S. Sanford, Z. Elisha-Feil, and D. Levens, The functional response of upstream DNA to dynamic supercoiling in vivo, Nat. Struct. Mol. Biol. 15, 146 (2008).

[32] S. S. Teves and S. Henikoff, Transcription-generated torsional stress destabilizes nucleosomes, Nat. Struct. Mol. Biol. 21, 88 (2014).

[33] A. Kaczmarczyk, H. Meng, O. Ordu, J. van Noort, and N. H. Dekker, Chromatin fibers stabilize nucleosomes under torsional stress, Nat. Commun. 11, 1 (2020).

[34] C. Naughton, N. Avlonitis, S. Corless, J. G. Prendergast, I. K. Mati, P. P. Eijk, S. L. Cockroft, M. Bradley, B. Ylstra, and N. Gilbert, Transcription forms and remodels supercoiling domains unfolding large-scale chromatin structures, Nat. Struct. Mol. Biol. 20, 387 (2013).

[35] R. Milo, P. Jorgensen, U. Moran, G. Weber, and M. Springer, Bionumbers—the database of key numbers in molecular and cell biology, Nucleic Acids Res. 38, D750 (2010).

[36] D. T. Gillespie, Exact stochastic simulation of coupled chemical reactions, J. Phys. Chem. 81, 2340 (1977).

